# Pesticidal Effects Of *Gmelina arborea* (Roxb) Powders On Beans Weevil (*Callosobruchus maculatus*) At Vom

**DOI:** 10.1101/2023.04.28.538680

**Authors:** Ojuolape Damilola Olanrewaju, Precious Josiah Edamaku, Ofodile Peter Nwite, Umoru Usman, Mohammed Murtala Aliyu

**Affiliations:** Department of Pest Management Technology, Federal College of Animal Health and Production Technology, Vom, Plateau State

**Keywords:** Pesticidal, *Callosobruchus maculatus*, *Gmelina arborea*, Mortality

## Abstract

Cowpea (Vigna unguiculata (L.) Walp. is one of the most nutritious grain legumes where it is valuable as a source of dietary protein yet most of the yields are lost as a result of infestation by pest especially Callosobruchus maculatus. cowpea weevil which is one of the most prevalent and a major destructive insect pest of stored legumes. Controlling of the pest by use of synthetic pesticides is raising serious concern on the environmental safety and consumer health hazards. This study was aimed at determining the pesticidal efficacy of Gmelina arborea stem bark and leaf powder on C. maculatus. The experiment was laid in a Completely Randomized Design (CRD) with three application rates (10g/200g grains, 10g/200g grains and 5+5g/200g cowpea grain) of the botanicals (leaf and stem bark powders and combination of leaf stem bark powders respectively) replicated three times was used in the assessment. The parameters and data collected in the experiment are; percentage of weevil mortality, Grain damaged, Number of Exit holes, newly emerged weevil, percentage of weight damaged. All data generated were statistically analyzed. The results of this study demonstrated the active potentials of these G. arborea plant parts as plant-derived pesticides against cowpea weevil. The effect of treated plant parts on percentage mortality rate showed significant difference (P < 0.05) over the untreated (control) as mixed leaf and stem bark powders recorded the highest mortality rate (85.00%). The leaf (51.70%) and stem bark (43.30%) powders also showed higher effect (51.70%) respectively while the lowest mortality rate was observed in the control (21.67%). The untreated (control) gave the highest number of newly emerged weevils (2.33) whereas the lowest (5.33) was found in Gmelina leaf + stem bark powders Gmelina leaf + stem bark powders attained the highest beans damage (11.00) while the lowest (9.67) was recorded in Gmelina leaf powders. The exit holes made by the weevils at the end of the experiment amongst G. arborea plant parts were highest (9.33) in Gmelina stem bark powders whereas lowest was recorded on Gmelina leaf + stem bark powders (5.33). Leaf + Stem bark powders was the most effective throughout the experiment followed by the leaf and stem bark powders respectively. In view of these findings G. arborea leaf and stem bark powders have a strong bioactivity and is effective against C. maculatus. Therefore, since these plant parts have no any adverse effects on the grains and safe to the environment, they are recommended for future usage in storage grains to control of C. maculatus.

## INTRODUCTION

Leguminous cowpea (*Vigna unguiculata* [L.] Walp) is a significant source of nutritional protein, vitamins, and minerals. It is mostly farmed for its nutritious grain legumes (Ehlers and Hala, 1997). (Singh *et al*., 2003). It is mostly grown and consumed by subsistence farmers in the famine- and malnutrition-prone semi-arid and sub-humid regions of Africa (DeBoer, 2003). The best use of cowpea, *Vigna unguiculata* L. Walp., as a dietary supplement and a sizeable source of income for the most vulnerable groups in West Africa is jeopardized by its difficulty in preservation (Akinkurolere *et al*., 2006; Ouédraogo, 2003). For many poor people in Nigeria, it is a significant cash and food crop because it is a significant and essential part of their diets. In Africa, where meat and other sources of animal protein are quite expensive, cowpea is comparatively inexpensive and helps many families meet their needs for protein (Nta *et al*., 2013). According to Lowenberg-Deboer and Ibro (2008), Nigeria is the world’s top producer and consumer of cowpeas. The FAO estimates that 3.3 million tons of dried cowpea grains were produced in 2000. Despite the cowpea’s numerous economic benefits and its relative significance to Nigeria’s economic development, it is unable to satisfy the population’s qualitative and quantitative needs. The vulnerability of cowpea to storage pests, primarily insects pests, is a significant hindrance (Tripathy *et al*., 2001).

Due to pod or seed infestation by bruchids, of which *Callosobruchus maculatus* fab. is the main pest, cowpea post-harvest storage is limited. (Sanon *et al*., 2005; Adedire *et al*., 2011). In Northern Nigeria and Northern Ghana, post-harvest losses of cowpea 3–4 months in storage due to *C. maculatus* infestation have been reported as high as 50% and 60%, respectively (Tanzubil, 1991). In Sub-Saharan Africa in general and Nigeria in particular, the cowpea weevil has been reported to be the most destructive pest of legume seeds. Its larva infest grains such cowpea, chickpea, Bambara groundnut, green gram, lentil, broad bean, and green pea. (Bagheri, 1996; Booth *et al*., 1990;).

Synthetic chemicals have been successful in controlling *Callosobruchus maculatus* in retailers. Many issues have arisen as a result of the extensive usage of these compounds, including insecticide resistance, consumer health risks, and environmental damage. Due to these issues, synthetic insecticides had to be replaced with natural substances that are safer for consumers and the environment, cheaper, and more effective at preventing insect infestations in stored grains. Thus, alternatives such as plant components from *Gmelina arborea* are needed to manage *C. maculatus*.

## MATERIALS AND METHODS

### EXPERIMENTAL SITE

The study was carried out in the Pest control laboratory of the Federal College of Animal Health and Production Technology Vom at the Chaha Campus.

### EXPERIMENTAL MATERIALS

The following materials were used for this study; *Vigna unguiculata* (Cowpea), *Gmelina arborea* (Leaf and Stem bark), Selo tape, Storage containers, Muslin cloth, Marker, Rubber bands, Hand lens, Weighing scale, Scissors

### EXPERIMENTAL PROCEDURE

#### Collection and Source of Cowpea grains

The cowpea was sourced from Bukuru market. Jos south L.G.A of plateau state. To kill any hidden infestation, the grains were properly sieved, hand-picked, and disinfected by being kept at

−5° C for seven days (Kophlar, 2003).

#### Weevil selection

100 pairs of adult cowpea weevils, *C. maculatus*, were placed into a 2 kg storage container holding 600 g of cowpea grains that were purchased at a nearby market in Bukuru, Jos Plateau state, Nigeria. The weevil colony was kept at a constant insectarium temperature with a relative humidity of 28.2°C and 75.5%.

In the pest management lab of the FCAH&PT, Vom, *C. maculatus* identification and sexing were done. Adults were then classified as male or female based on the rostrum’s length (the female has a comparatively longer rostrum than the male).

#### Collection and preparation of plant powders

The plant components of *Gmelina arborea* were taken immediately from the tree and brought to the Central Diagnostic Laboratory, National Veterinary Research Institute, Vom. Both plant sections were cleaned with clean tap water after being dusted to eliminate dirt. In the laboratory, they were first naturally air dried. Later, a mechanical grinding device was used to independently crush the dried components into fine powder. The powders were then divided into distinct plastic bags and kept in storage to maintain their high quality before use.

10 males and 10 females of the adult bean weevil (*C. maculatus*) were introduced to 200g of cowpea grains in a storage container. In the containers designated for their treatment, 10g of *G. arborea* Leaf and Stem Bark Powders were measured and carefully combined with the grains. To keep the insects from escaping, a transparent muslin cloth was placed over each box. Within a 2-week period, all parameters were measured at regular intervals. After an infestation of two weeks, the setup was examined, and dead adults were tallied. The Asawalam *et al*. formula was used to determine the adult mortality rate (2006).

### EXPERIMENTAL DESIGN

A 4 × 3 factorial experiment laid in a Complete Randomized Design (CRD) which was replicated 3 times

### DATA COLLECTION

Data were collected by checking for the following parameters; Mortality rate, Number of exit holes, newly emerged weevil, Initial and final weight of the cowpea grains and Level of beans damage.

#### Mortality test

With few modifications, the mortality test was carried out using the same methods as Chebet *et al*. (2013). Twenty (20) adult (male and female) G. arborea plant stem bark and leaf powders were combined with 200g of cowpea grains at various dosages. After one week and two weeks, the number of dead adults was counted, accordingly. The adult mortality rate was calculated according to the formula of Asawalam *et al*. (2006).

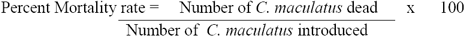

### STATISTICAL ANALYSIS

Data were subjected to analysis of variance (ANOVA) using the statistical package SPSS 23.0 software (SPSS, 2017). Means were separated using Least Significant Difference (LSD).

## RESULTS

The effectiveness of the various plant parts of *Gmelina arborea* on *Callosobruchus maculatus* on the weevil mortality is significantly different (P **≤** 0.05) throughout all weeks, as shown in Table 1. Week one mortality was 8.33% for the control group. The maximum and lowest mortalities were noted for the *G. arborea* plant sections on leaf + stem bark powders (35.00%) and stem bark powder (21.67%), respectively. Leaf + stem bark powders had the greatest impact during week two (85.00%). Powders of the stem bark (43.30%) and leaf (51.70%) also had a stronger impact, respectively. The control group had the lowest mortality rate (21.67%).

**Table 1:** Efficacy of *Gmelina aborea* Leaf and Stem bark on the Mortality rate of *C. maculatus*.

| Treatment | WEEKS |  |
| --- | --- | --- |
|  | ONE | TWO |
| Control (Untreated) | 8.33<sup>c</sup> | 21.67<sup>b</sup> |
| *Gmelina leaf* (10g) | 10.00<sup>c</sup> | 51.70<sup>ab</sup> |
| *Gmelina stem bark* (10g) | 21.67<sup>b</sup> | 43.30<sup>ab</sup> |
| *Gmelina leaf* + *stem bark* (5+5g) | 35.00<sup>a</sup> | 85.00<sup>a</sup> |
| P value | 0.000 | 0.024 |

**Legend: Means that do not share a superscript within the same column are significantly different from each other at ( $P \leq 0.05$ )**

According to Table 2, there were variations in the population of freshly emerged weevils during the experiment that were statistically significant (P **≤** 0.05). Compared to other plant components, the effect of *Gmelina* leaf powders had a significant impact on the weevil population (P 0.05), as it recorded the highest (2.67). The control (untreated) group recorded the greatest number of freshly emerging weevils at the conclusion of the first week (6.33) *Gmelina* leaf + stem bark powders had the lowest newly emerged weevils in week two (5.33) and the highest number (2.33) (control). Both the leaf (8.67) and stem (9.33) powders of *Gmelina* included a lot of newly emerged weevils. Table 3 shows the effect of different *G. arborea* plant parts on the number of exit holes. At week 1, there was no significant difference (P < 0.05) in the number of exit holes between *Gmelina* plant parts. Control had the highest number (8.00) of exit holes while *Gmelina leaf + stem bark* powders had the lowest number (4.33) of exit holes. The exit holes made by the weevils at the end of the experiment (week 2) amongst *G. arborea* plant parts were highest (9.33) in *Gmelina stem bark* powders whereas lowest was recorded on *Gmelina leaf + stem bark* powders (5.33). Control had the highest number (15.00) of exit holes at the end of week 2

**Table 2:** Effects of *G. aborea* Leaf and Stem bark on Newly emerged weevils of *C. maculatus*.

| Treatment | WEEKS |  |
| --- | --- | --- |
|  | ONE | TWO |
| Control (Untreated) | 6.33<sup>a</sup> | 15.00<sup>b</sup> |
| *Gmelina* leaf (10g) | 2.67<sup>ab</sup> | 8.67<sup>ab</sup> |
| *Gmelina* stem bark (10g) | 2.00<sup>b</sup> | 9.33<sup>ab</sup> |
| *Gmelina* leaf + stem bark (5+5g) | 1.33<sup>b</sup> | 5.33<sup>a</sup> |
| P value | 0.010 | 0.049 |

**Legend: Means that do not share a superscript within the same column are significantly different from each other at ( $P \leq 0.05$ )**

**Table 3:**
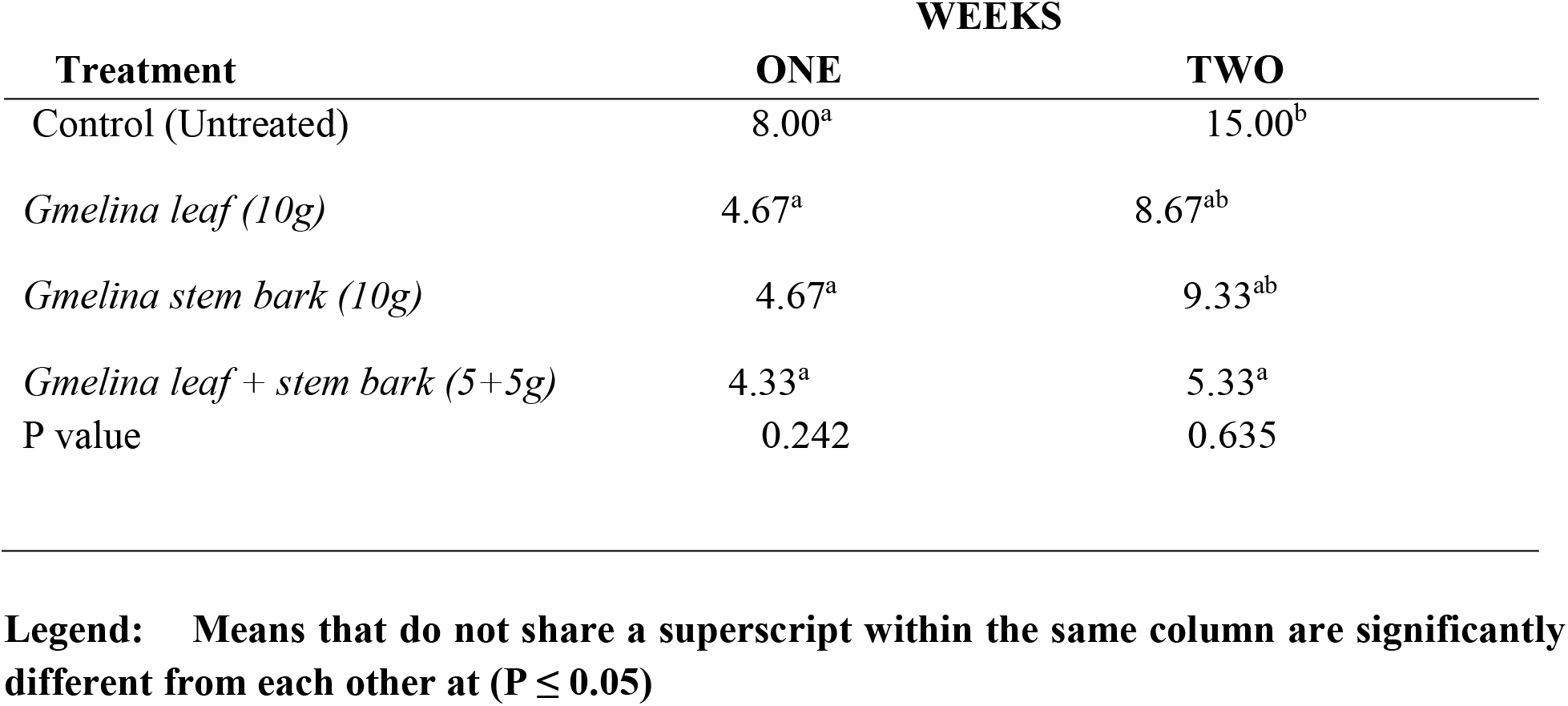
Effects of *G. aborea* Leaf and Stem bark on Number of Exit Holes on beans grains

Table 4 shows the effect of different *G. arborea* plant parts on the number of beans damaged. The number of beans damaged in each *Gmelina* plant part was the same throughout the trial, which was significant (P≤ 0.05). At week 1, the highest number of beans damaged was observed in *Gmelina leaf* powders (5.33) while *Gmelina leaf + stem bark* powders had the lowest number of damaged beans (2.33). Among *G. arborea* plant parts at week 2, *Gmelina leaf + stem bark* powders attained the highest beans damage (11.00) while the lowest (9.67) was recorded in *Gmelina leaf* powders.

**Table 4:**
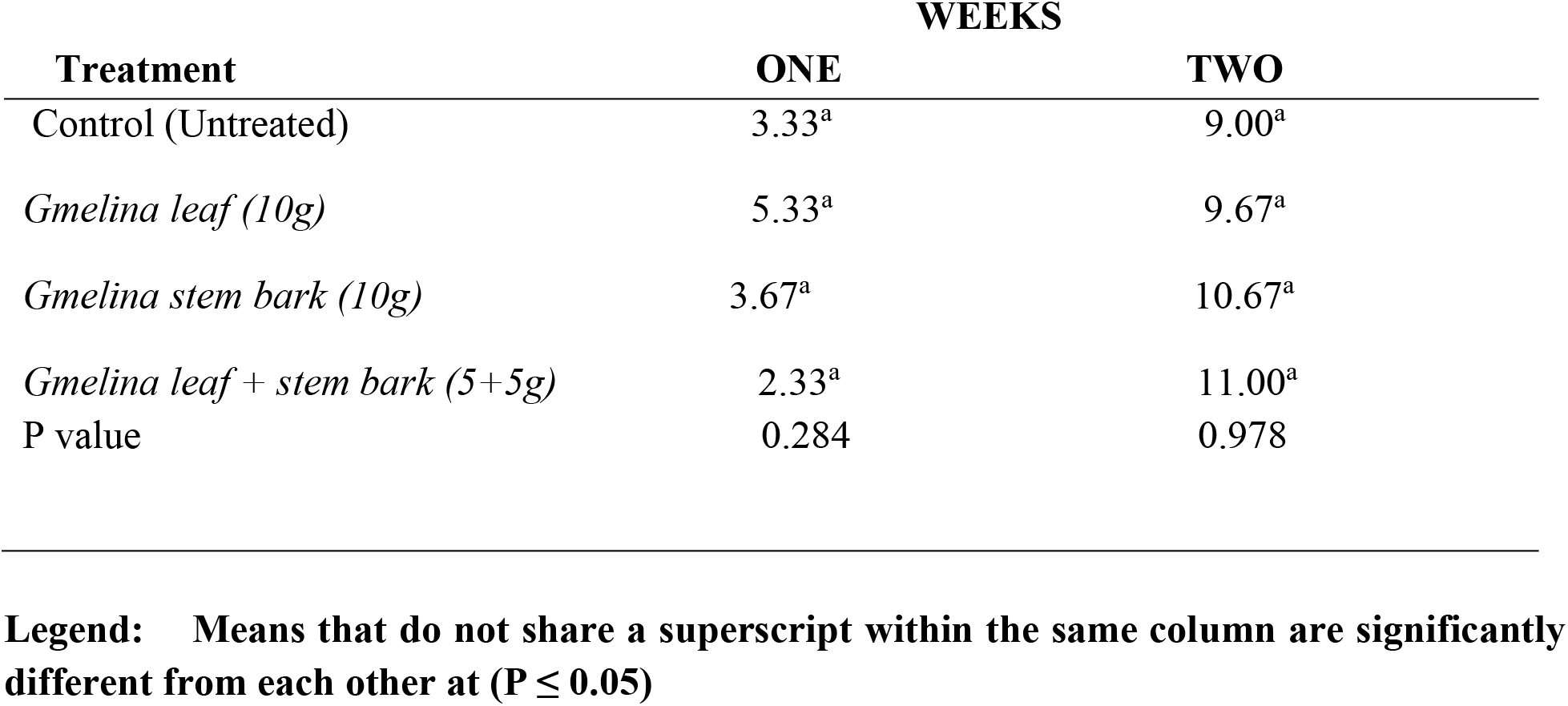
Efficacy of *G. aborea* Leaf and Stem bark on the number of beans damaged

| Treatment | WEEKS |  |
| --- | --- | --- |
|  | ONE | TWO |
| Control (Untreated) | 3.33<sup>a</sup> | 9.00<sup>a</sup> |
| *Gmelina leaf* (10g) | 5.33<sup>a</sup> | 9.67<sup>a</sup> |
| *Gmelina stem bark* (10g) | 3.67<sup>a</sup> | 10.67<sup>a</sup> |
| *Gmelina leaf + stem bark* (5+5g) | 2.33<sup>a</sup> | 11.00<sup>a</sup> |
| P value | 0.284 | 0.978 |

**Legend: Means that do not share a superscript within the same column are significantly different from each other at ( $P \leq 0.05$ )**

A significant difference in the weight loss of grains following the experiment is shown in Table 5. Among the plant parts of *G. arborea, Gmelina* leaf + stem bark powders had the lowest weight loss (140.00), while *Gmelina* leaf powders had the largest weight loss (172.00). Untreated (control) recorded 185.00.

**Table 5:**
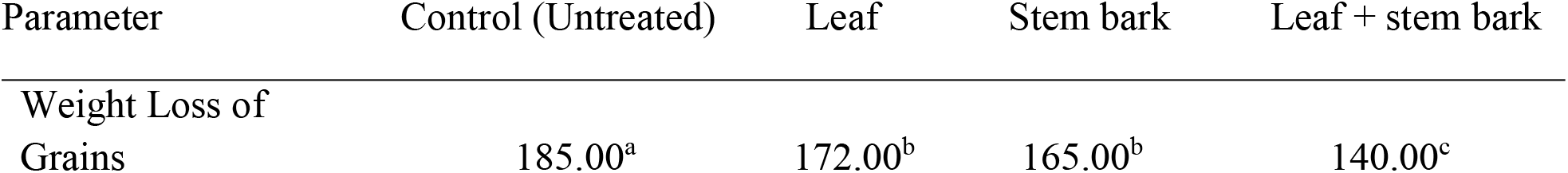
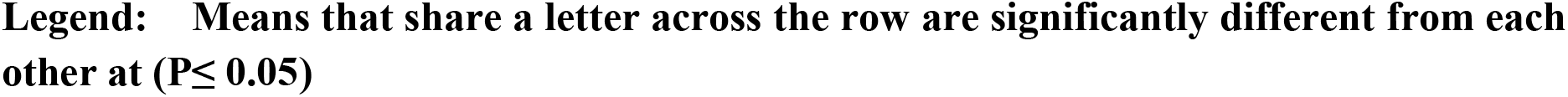
Effect of *G. aborea* Leaf and Stem bark on the Weight loss of grains after weeks

| Parameter | Control (Untreated) | Leaf | Stem bark | Leaf + stem bark |
| --- | --- | --- | --- | --- |
| Weight Loss of Grains | 185.00<sup>a</sup> | 172.00<sup>b</sup> | 165.00<sup>b</sup> | 140.00<sup>c</sup> |

**Legend: Means that share a letter across the row are significantly different from each other at ( $P \leq 0.05$ )**

## DISCUSSION

Despite the fact that the leaves and stem bark utilized in this study came from the same plant, the results, as shown in Tables 1, 2, and 3, showed that they had different effects on adult C. maculatus. According to the percentage mortality results shown in Table 1, Stem bark + leaf powders at a concentration of 10g (5g+5g) had the highest percentage mortality of 85.00%, contradicting Oladejo et al. (2020), who found that the combination of stem bark and leaf powders had the least impact on weevil mortality. The stem bark powder, which had a percentage mortality of 43.30% at a concentration of 10g, had the least impact. With increasing exposure time, it was observed that the mortality rate of *C. maculatus* caused by plant parts increased. Thus, the study’s results revealed that when days increased from week one to week two, the mean values obtained for each treatment increased as well. This is feasible due to the fact that active components of *G. arborea* require greater concentration and longer amount of time to bio-magnify in *C. maculatus*. This is consistent with research conducted by Kemabonta and Falodu (2013) on the effectiveness of three plant products as post-harvest grain protectants against *Sitophilus oryzea* L (Coleoptera: Lurculionidae) on stored wheat (*Triticum aestivum*). They found that *S. oryzea* mortality is based on the exposure time and concentration of the plants used. The untreated (control) grain provides a liberated environment where weevils can proliferate without hindrance, which accounts for the highest feeding rates (Dolob, 2002). Moreover, the plant has been reported to possess anti-microbial, anti-feedant, and repellent properties (Hammuel *et al*., 2011). The obstruction of insect spiracles by dust particles from pulverized plant powder results in insect mortality (Fernando and Karunaratne, 2012). According to a different study by Kedia *et al*. (2015), botanicals can also enter an insect’s body through the respiratory system and kill it. The effect is dose dependant, and combining the various plant parts did not improve insect newly emerged. This is comparable to the work by Oladejo *et al*. (2020), which revealed that the combination of stem bark and leaf powders had only marginal success in suppressing *S. zeamais*. Due to the adult *C. maculatus’* inability to feed, *G. arborea* might be found to exhibit the capacity to act as smothering material with the potential to impede respiration. This is in accordance with studies by Udo (2011), who discovered that plant oils can suffocate *C. maculatus*. The impact of various plant materials on insects may vary depending on a number of variables, including chemical composition and species sensitivity (Aktar, 2004). Following the experiment, there were significant differences in the weight loss of grains (P 0.05). (Table 5). The highest weight reduction was seen in the untreated (control) group, which differs noticeably from all other treatments (table 5). In contrast to all the previous treatments, this was brought on by weevil damage to grains. This is consistent with Duke (2001), who reported that weevils can cause grain to lose 80–100% of its weight if it is not treated for a long time. The results of this study also showed that the weight losses brought about by *C. maculatus* on cowpea grains treated with various dosages of *G. arborea* Leaf, Stem bark, and leaf + stem bark powders were less than those brought about on the control (untreated), which may be due to the anti-feeding qualities of the *G. arborea* Leaf, Stem bark, and leaf + stem bark powders, as well as the insecticidal activities of the phytochemical components (Chothani and Patel, 2018). This outcome corresponds with Idoko and Adebayo’s (2011) observation that the control group experienced the greatest weight reduction from *D. porcellus* on yam chips (untreated) while yam chips treated with *G. arborea* Leaf, Stem, and Bark Powders had the lowest treatment. The research of Angaye *et al*. (2017), which showed the effectiveness of leaf extracts of *Gmelina arborea* against mosquito larvae, is similar to the findings in this study. By combining the powders of *Bridelia ferruginea (*Benth), *Blighia sapida* (Juss), and *Khaya senegalensis* (Cronquist), Estelle *et al*. (2018) demonstrated the effectiveness of the mixtures against *Dinoderus porcellus* that infests yam chips.

Iswarya *et al*. (2017) report that *G. arborea* contains substances with pharmacological properties. Strong antibacterial properties of *G. arborea* plant parts were also discovered by studies by De Bruyne *et al*. (1999) and Kaswale *et al*. (2012). Furthermore, it has been said that the plant contains oil that is used to protect wood from termites, particularly gregarious ones (Kareu *et al*., 2010). As a result, the present research confirmed that *G. arborea* has pesticidal effects due to the presence of tanin, cardiac glycoside, and phenol glycoside (Ahmad *et al*., 2001).

## CONCLUSION

This research showed the pesticidal effectiveness of *G. arborea’s* leaf and stem bark. According to the findings of this study, Gmelina arborea is effective against *Callosobruchus maculatus*. However, the dose and plant parts affect efficacy. The results of this study demonstrated that *C. maculatus* may be controlled and prevented by using the leaf and stem bark powder of *G. arborea* at different dosages. High mortality rate was seen in Leaf + Stem Bark Powder (5g+5g). Also, it was discovered that leaf and stem bark powder both had strong insecticidal effects on bean weevils. In contrast to synthetic chemical insecticides that pose risks to the environment’s health and expose people to lethal doses, stem bark powders proved to be the most effective throughout the experiment, followed by leaf and stem bark powders, respectively.

## RECOMMENDATION

Since the plant’s eco-friendliness and accessibility, it should be encouraged that poor resource farmers and food vendors utilize powdered *Gmelina arborea* leaf and stem bark to control the beans weevil in stored cowpea grains. This will decrease the application of synthetic insecticides, reducing the chances of contact and usage-related hazards.

The fruit and root of *G. arborea*, among other parts, could also be examined in different forms. To conduct this research and see if better results might be reached, several *G. arborea* components could be combined. Also, more study should be conducted to identify other insecticidal active components that can be used in the formulation of plant-based pesticides. It is also encouraged to conduct more research to find out how well this botanical works against other cowpea pests.

